# Protein kinase C inhibitor suppresses 2-cell stage development and perinuclear vesicle formation in mouse zygotes

**DOI:** 10.1101/2025.06.23.661161

**Authors:** Toru Suzuki, Yuriko Sakamaki

## Abstract

Nuclear structure and nucleocytoplasmic interaction are closely linked to the regulation of cellular and higher-order biological processes. An alternative pathway for nucleocytoplasmic transport involving perinuclear vesicle formation and release, termed nuclear envelope budding (NEB), has been observed in diverse species and cell types. NEB-like events have also been reported in mammalian zygotes, including mice. However, their molecular basis remains unclear. Recent results suggest that protein kinase C (PKC) signaling regulates perinuclear vesicle biogenesis during NEB. While multiple PKC isoforms are expressed in mouse oocytes, their functions in zygotes are not fully understood. We investigated the effects of pharmacological PKC inhibition on zygotic development and NEB-like perinuclear vesicle formation. Our results suggest that NEB-like events in mouse zygotes may involve PKC-dependent mechanisms and that PKC activity might be critical for the 1-to 2-cell transition.

## 1. Introduction

The nucleus is closely involved in regulating cellular and higher-order biological processes. Its structure, function, and nucleocytoplasmic communication are controlled by intracellular molecular signals. Although the nucleus and cytoplasm are separated by the nuclear envelope, cells possess regulated mechanisms enabling material exchange between them [1]. Although the primary pathway of nucleocytoplasmic transport involves the nuclear pore complex (NPC), an alternative route, nuclear envelope budding (NEB), has been identified [2]. NEB entails the encapsulation of nuclear contents by the inner nuclear membrane, formation of vesicular structures within the perinuclear space, and release into the cytoplasm. It is associated with protein quality control, signal transduction, and transport of large nuclear cargo [2,3] and may contribute to organismal homeostasis during aging [4].

NEB-like vesiculation has been observed in mammalian zygotes and early preimplantation embryos including mice [5–7]. However, its molecular nature remains unclear. Researchers described electron-dense, non-basic material enclosed by the inner nuclear membrane of pronuclei and vesicle budding into the cytosol, suggesting a noncanonical nuclear export route in early development. We refer to this process as “early embryonic NEB (eeNEB)” (see Discussion) [8].

Although the molecular basis of NEB remains unclear, studies in Drosophila [2] suggest that specific signaling pathways, including atypical protein kinase C (aPKC), are involved in its regulation [8]. aPKC appears to initiate perinuclear vesicle formation. Additionally, NEB-like nuclear egress of viral capsids reportedly requires PKC activity [8]. In contrast to classical (cPKC) and novel (nPKC) isoforms, which respond to Ca^2+^ and diacylglycerol (DAG), aPKCs are insensitive to both [9].

In mice and other species, sperm-induced Ca^2^+ signaling plays an essential role in fertilization and oocyte activation by promoting meiotic resumption and initiating embryogenesis [10]. One prevailing model suggests that sperm-derived phospholipase C zeta (PLCζ) hydrolyzes phosphatidylinositol 4,5-bisphosphate (PIP_2_) to generate inositol 1,4,5-trisphosphate (IP_3_), and possibly DAG, thereby triggering Ca^2+^ release via IP_3_-sensitive intracellular channels.

Several PKC isoforms are expressed at the mRNA and protein levels in mouse oocytes and zygotes [11–14]. Although various PKCs have been linked to fertilization and preimplantation development [15–21], recent studies (e.g., [22]) showed that oocyte activation and even full-term development can occur without calcium transients or sperm-derived factors, questioning the essential roles of calcium- and DAG-dependent PKCs in zygotes. In this study, we examined PKC function during early embryogenesis and its role in regulating eeNEB using pharmacological inhibitors.

## 2. Materials and methods

### 2.1. Animals

All animal experiments were approved by the Institutional Animal Care and Use Committee of Institute of Science Tokyo. B6D2F1 mice were obtained from CLEA Japan and maintained on a 12-hour light/dark cycle. Food and water were provided ad libitum to the mice.

### 2.2. Chemical inhibitors

PKCι inhibitor 1 (compound 19, iPKCι, MCE), Go 6983 (Funakoshi, Japan), and Sotrastaurin (Sigma-Aldrich, USA) were dissolved in dimethyl sulfoxide (DMSO; 10–50 mM stocks) and stored at –20 °C. For experiments, inhibitors were diluted in potassium simplex optimization medium (KSOM) to final concentrations as indicated. iPKCι inhibits PKCι(λ), α, ε, and ζ with IC_50_s of 0.34, 0.34, 1.6, and 0.51 µM, respectively [23]. Go 6983 targets PKCα, β, γ, δ, and ζ (IC_50_s: 7, 7, 6, 10, and 60 nM, respectively) [24]. Sotrastaurin inhibits PKCθ, β, α, η, δ, and ε with K_is_ of 0.22, 0.64, 0.95, 1.8, 2.1, and 3.2 nM, respectively [25].

### 2.3. Oocyte collection, in vitro fertilization and embryo culture

Female B6D2F1 mice (3–4 weeks old) were superovulated via intraperitoneal injection of 0.1 Ml HyperOva (Kyudo, Japan) [26] and 7.5 IU hCG (Asuka Animal Health, Japan) 48 h apart. Fourteen to fifteen hours post-hCG, mice were euthanized and cumulus–oocyte complexes (COCs) were retrieved from oviducts into Human tubal fluid (HTF) medium (Kyudo, Japan). Prior to this, 8–12-week-old B6D2F1 males were sacrificed and sperm were collected from the cauda epididymis by 1-hour swim-up in HTF. Preincubated sperm were added to COCs in HTF for *in vitro* fertilization (IVF) and co-incubated for 2.5–6 h. Fertilized oocytes, identified by second polar body or pronuclei, were washed and cultured in KSOM (ARK Resource, Japan) under 5% CO_2_at 37 °C. Zygotes were exposed to test compounds in KSOM for up to 5 days.

Germinal vesicle (GV)-stage COCs were collected as described [27], with minor changes. Female mice received 0.1 mL HyperOva; 48 h later, ovaries were excised and minced in M2 + 150 µM 1-Methyl-3-Isobutylxanthine (IBMX; Nacalai, Japan). Cumulus cells were removed by pipetting and GV oocytes were either fixed or imaged.

### 2.4. Oocyte activation by N,N,N′,N′-Tetrakis(2-pyridylmethyl)ethylenediamine

Oocyte activation by N,N,N′,N′-Tetrakis(2-pyridylmethyl)ethylenediamine (TPEN) was performed as previously described [22], with minor modifications. Unfertilized oocytes were collected in KSOM and incubated with 100 µM TPEN for 1 h. Activated oocytes were cultured in KSOM with 100 µM ZnSO_4_ for 3 h, following a 1-hour TPEN washout. Subsequently, 1-cell embryos were transferred to KSOM without ZnSO_4_ and cultured for 4 h. Nine hours after TPEN treatment began, 1-cell embryos were either photographed or fixed for electron microscopy.

### 2.2. Electron microscopy

GV oocytes, zygotes cultured with or without PKC inhibitors, and TPEN-activated 1-cell embryos were fixed in 2.5% glutaraldehyde in 0.1 M phosphate buffer (PB) for 2 h at 4 °C, washed in 0.1 M PB containing 0.1% Bovine serum albumin (BSA; Sigma-Aldrich, USA), and post-fixed in 1% osmium tetroxide (OsO4) buffered with 0.1 M PB for 2 h. After washing with PB, cells were resuspended in 2% gelatin (Sigma-Aldrich, USA) and pelleted. Microcentrifuge tubes were plunged into ice-cold water to quickly solidify the gelatin with the cells. The tip of the tube was cut open and the cell pellets were cut into 1-mm^3^ blocks. Blocks were dehydrated in a graded series of ethanol and embedded in Epon 812. Ultrathin sections (70 nm thick) were mounted on a silicon wafer, double-stained with uranyl acetate and lead citrate, and examined using a scanning electron microscope (SEM, JSM-7900F, JEOL, Tokyo, Japan). Electron micrographs were acquired at magnifications ranging from 1,000× to 30,000×.

### 2.6. Image analysis

Images of oocytes and embryos were captured using an inverted microscope (IX73, Olympus, Japan) or SEM and analyzed with ImageJ software. The pronuclear perimeter was estimated using major and minor axes and Ramanujan’s ellipse circumference approximation [28]. The perinuclear vesicle area was calculated from their major and minor axes.

### 2.7. Statistics

Developmental outcomes among multiple groups were analyzed using Fisher’s exact test with Holm correction, with a statistical significance of *p* < 0.05. To assess the effects of PKC inhibition, perinuclear vesicle counts per zygote were normalized to pronuclear perimeter length (expressed as vesicles/µm), allowing the comparison between PKC inhibitor-treated and control groups. Similarly, the total perinuclear vesicle area was normalized to the pronuclear perimeter (area/µm) for each zygote. Comparisons between two groups were conducted using two-tailed Student’s t-tests, with a significance of *p* < 0.05. All data were derived from oocytes or embryos obtained in two independent experiments.

## 3. Results

### 3.1. Effects of PKC inhibitors on mouse preimplantation embryogenesis

Three hours after IVF initiation (Fig. 1A), zygotes were cultured in KSOM medium supplemented with 3.3–100 µM iPKCι, Go 6983, and sotrastaurin or without inhibitor (Figs. 1B–E, S1,

**Fig. 1.**
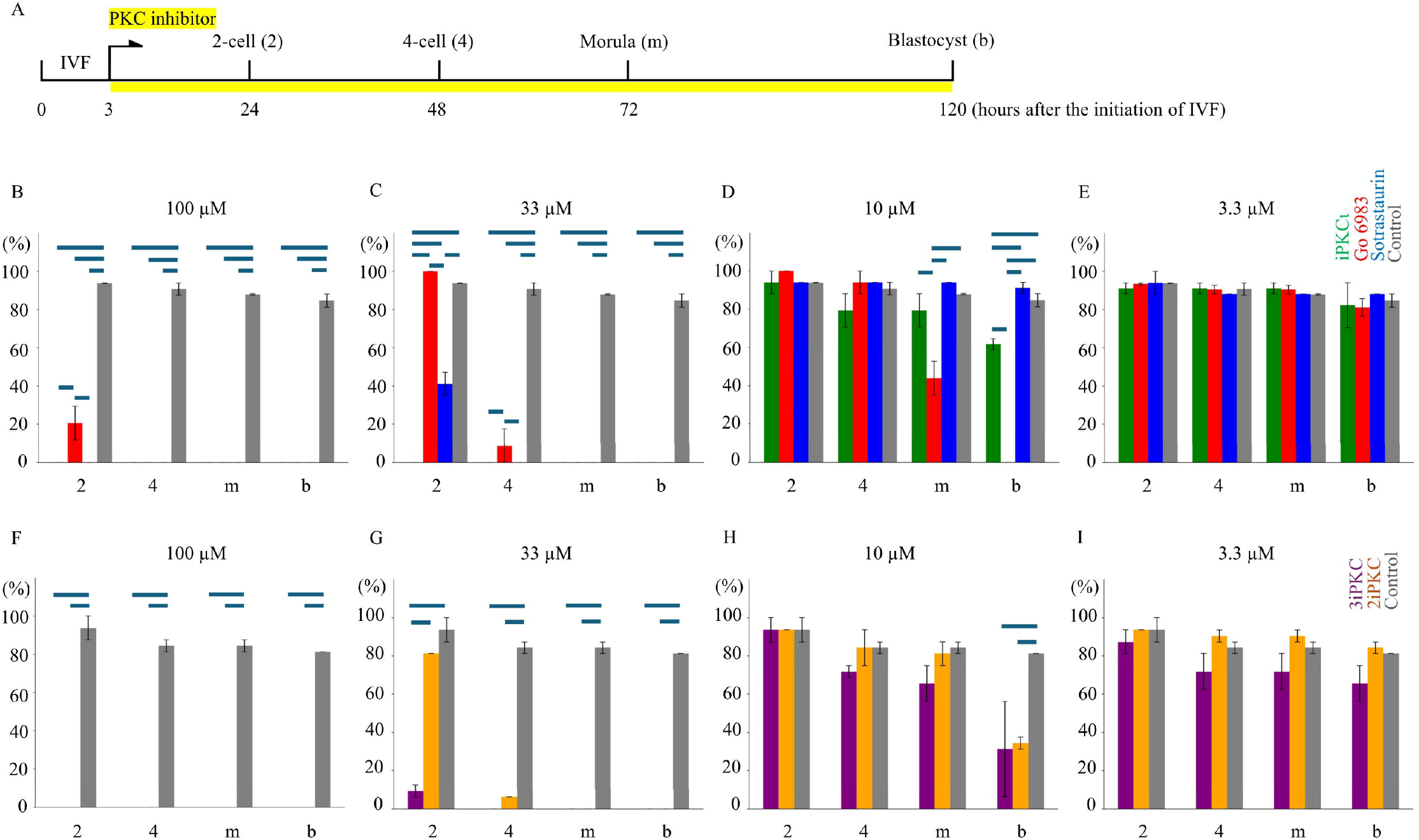
Effect of PKC inhibitors on the embryonic development of mouse zygotes. (A) Schematic timeline of *in vitro*-fertilized (IVF) embryo culture with protein kinase C (PKC) inhibitors. Embryos were cultured with inhibitors from 3 to 120 h after IVF initiation (yellow shading). The development was assessed in the 2-cell (2), 4-cell (4), morula (m), and blastocyst (b) stages. Average developmental rates ± standard deviation (SD) relative to total zygotes were calculated. (B–E) Preimplantation development with individual PKC inhibitors: iPKCι (green), Go 6983 (red), sotrastaurin (blue), at 3.3–100 µM. Untreated controls (gray) are shown for comparison (data from two independent experiments). (F–I) Preimplantation development with combined PKC inhibitors: Go 6983 + sotrastaurin (2iPKC, orange) or iPKCι + Go 6983 + sotrastaurin (3iPKC, purple) at total concentrations ranging from 3.3–100 µM. Controls (gray) are shown. The lines above colored bars (B–I) show the significant difference (*p* < 0.05) between 2 groups. IVF: *in vitro* fertilization, iPKCι: PKC inhibitor 1.

Tables S1–2). At 100 µM, all pairwise comparisons for 2-cell development showed significant differences, except for the iPKCι and sotrastaurin groups. At 33 µM, all comparisons showed significant differences, except for the Go 6983 vs. control comparison. Notably, zygotes treated with iPKCι showed 0% development to the 2-cell stage while maintaining a morphologically usual 1-cell appearance. Concentrations of ≤10 µM did not significantly affect 2-cell development in any group. At 10 µM, both morula and blastocyst formation were significantly reduced in the Go 6983 group compared with the other groups. Blastocyst development in the iPKCι group also differed significantly from that in other groups.

Zygotes were then treated with dual (Go 6983 + sotrastaurin; 2iPKC) or triple (iPKCι + Go 6983 + sotrastaurin; 3iPKC) inhibitor mixtures at total concentrations ranging from 3.3–100 µM (Figs. 1F–I, S2, Tables S3–4). At ≤33 µM (≤16.5 µM each) of 2iPKC, 81.3%–93.8% developed to 2-cell within 24 h. At 100 µM (50 µM each), all arrested in the 1-cell stage. Blastocyst development was significantly impaired at ≥10 µM compared with the control.

With 3iPKC, 9.4% of zygotes developed to 2-cell at 33 µM. Blastocyst development was significantly reduced at ≥10 µM compared with the control.

### 3.2. Perinuclear vesicles in mouse GV oocytes and early preimplantation embryo

Previously, perinuclear vesicles appearing as early as 6 h post-sperm penetration and persisting through late 1-cell and early 2-cell stages but absent in GV oocytes and mid to late 2-cell stages were reported [7]. To reassess this, electron microscopy was performed on GV oocytes, 3-h post-IVF zygotes (IVF 3 h), and 30-h 2-cell embryos (IVF 30 h). NEB-like vesicles were absent in GV oocytes (n = 14) and IVF 30 h embryos (n = 12; Figs. 2A, C, E, G, I, K, and S3). In the IVF 3 h group (n = 8), clear perinuclear vesicles were observed in one zygote containing a sperm tail cross section [29], with electron-dense perinuclear microprotrusions in others (Figs. 2B, F, J, and S3H-J). Ultrastructure, including a nuclear envelope lipid bilayer, cytoplasmic lattices [30], intranuclear annulate lamellae [7,31], and endolysosomal vesicular assemblies (ELVAs) [32] were observed (Figs. S3A–K).

**Fig. 2.**
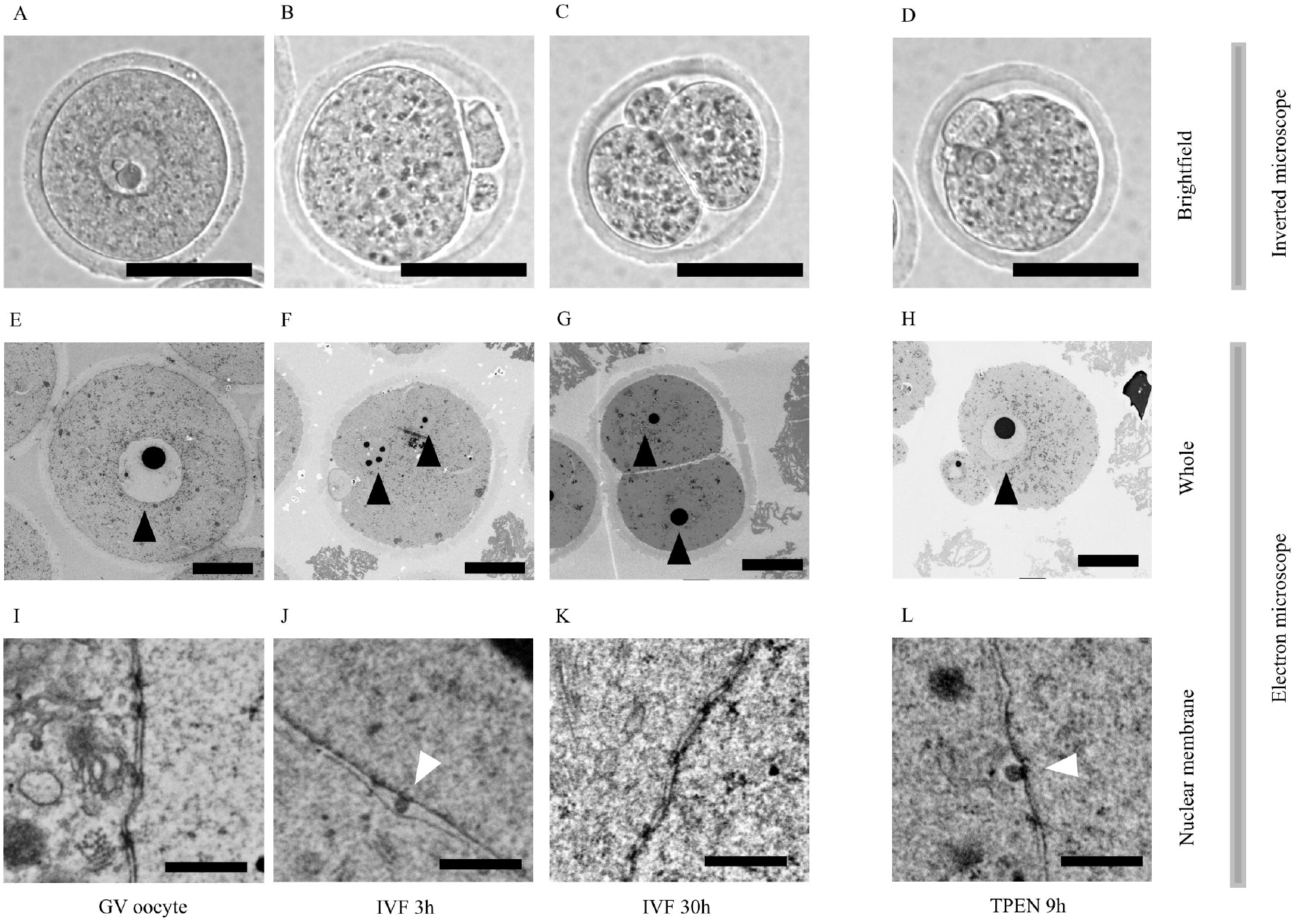
Electron microscopy and brightfield images of mouse oocytes and embryos. (A–D) Representative brightfield images (200×) of germinal vesicle (GV) oocytes (A), embryos 3 h (IVF 3h, B) and 30 h (IVF 30h, C) after IVF initiation and TPEN-activated embryos 9 h after the initiation of the treatment (TPEN 9h, D). Black scale bar = 50 µm. (E–H) Representative electron microscopy images (1,000×) of GV oocytes (E), IVF 3h (F), IVF 30h (G), and TPEN 9h embryos (H). Black arrowheads indicate nuclei. Scale bar = 20 µm. (I–L) Electron microscopy images (30,000×) of nuclear membranes in GV oocytes (I), IVF 3h (J), IVF 30h (K), and TPEN 9h (L). White arrowheads indicate perinuclear vesicles. Scale bar = 0.5 µm. IVF: *in vitro* fertilization, iPKCι: PKC inhibitor 1, TPEN: N,N,N′,N′-Tetrakis(2-pyridylmethyl)ethylenediamine.

### 3.3. Effects of PKC inhibitors on perinuclear vesicle formation of mouse zygote

Electron microscopy was performed on zygotes with or without PKC inhibitors. Without inhibitors, perinuclear vesicles were observed 9 h post-IVF with an average area of 9630.8 ± 312.7 nm^2^ (n = 258; Fig. 3).

**Fig. 3.**
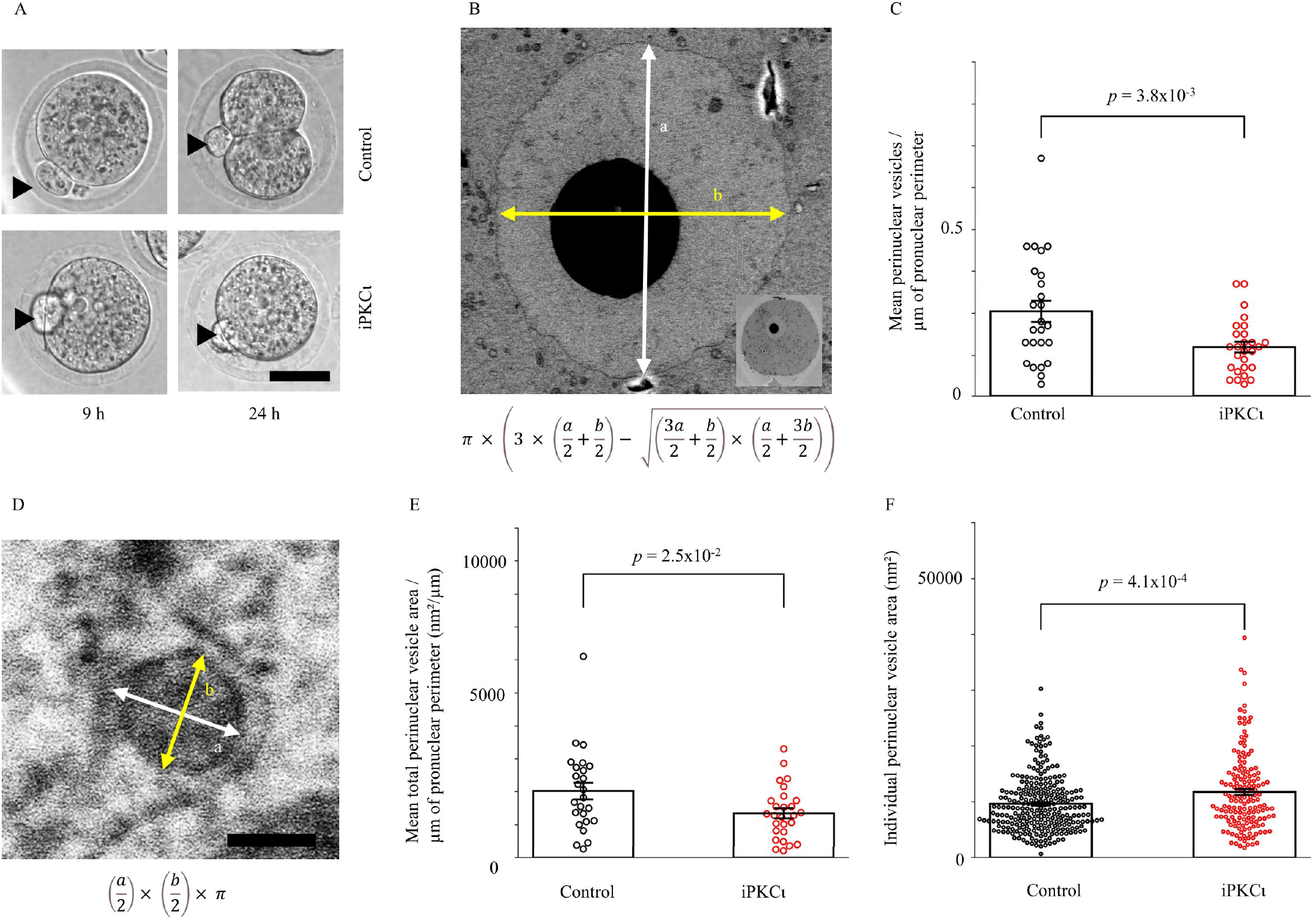
Effects of PKC inhibitor on perinuclear vesicle formation in mouse zygotes. (A) Representative brightfield images (200×) of embryos treated with 33 µM iPKCι or untreated controls at 9 and 24 h after IVF initiation. Zygotes treated with iPKCι arrest at the 1-cell stage at 24 h, whereas controls develop to the 2-cell stage. Black arrowheads mark second polar bodies. All images share the same scale; scale bar = 50 µm. (B) Electron microscopy image (1,000×) of a pronucleus from an iPKCι-treated zygote 9 h after IVF initiation, with an approximate formula for pronuclear perimeter calculation. The inset shows a full zygote view. Double-headed arrows indicate the major and minor axes of the vesicle. (C) Average number of perinuclear vesicles per unit pronuclear perimeter length. Values were averaged per zygote and then compared between treated (n = 25) and control (n = 27) groups. The statistical significance was assessed with a two-tailed Student’s t-test (*p* < 0.05). (D) Electron microscopy image (30,000×) of a perinuclear vesicle, with a formula for area calculation. Double-headed arrows indicate the major and minor axes of the vesicle. Scale bar = 100 µm. (E) Comparison of the average total perinuclear vesicle area per unit pronuclear perimeter length of iPKCι-treated (n = 25) and control (n = 27) groups. (F) Comparison of the size of individual perinuclear vesicles of iPKCι-treated (n = 258) and control (n = 153) groups. Statistical analysis was carried out via two-tailed Student’s t-tests (*p* < 0.05: significantly different). IVF: *in vitro* fertilization, iPKCι: PKC inhibitor 1.

Treatment with 33 µM iPKCι for 6 h starting 3 h after IVF initiation, which induces 1-cell arrest (Fig. 3A), retained vesicles but reduced their density along the pronuclear perimeter to 60% of the control (0.12 ± 0.01/µm vs. 0.20 ± 0.03/µm; *p* = 3.8×10^−3^; n = 27 and 25; Figs. 3B–C). The total vesicle area per µm pronuclear perimeter was also significantly lower with inhibitor (1349.7 ±152.2 vs. 2020.3 ± 251.9 nm^2^/µm; *p* = 0.024; n = 27 and 25; Figs. 3D–E). Conversely, the mean vesicle area was larger in treated zygotes (11751.1 ± 566.0 nm^2^; n = 153) than controls (*p* = 4.1×10^−4^; Fig. 3F).

At 100 µM 3iPKC, also causing 1-cell arrest (Fig. S4A), the vesicle density (0.03 ± 0.01/µm vs. 0.10 ± 0.02/µm; *p* = 1.9×10^−4^; n = 22 and 24, Figs. S4B–C) and total area (300.2 ± 56.7/µm vs. 944.2 ± 142.7/µm; *p* = 2.0×10-^4^; n = 22 and 24, Figs. S4B and D) of the treated group were markedly reduced compared with those of the controls. However, the mean vesicle area did not significantly differ from that of the control group (8266.7 ± 776.0 vs. 9370.2 ± 430.2 nm^2^; *p* = 0.22; n = 41 and 140, Fig. S4E).

### 3.4. Effects of PKC inhibitors on the ultrastructure of pronuclei in mouse zygotes

During perinuclear vesicle quantification, a higher electron density was observed on the nucleoplasmic side of the inner pronuclear membrane in PKC inhibitor-treated zygotes (Figs. 4A and S5–6). Electron microscopy was used to quantify the relative signal intensity at the inner and outer nuclear membranes in zygotes treated with 33 µM iPKCι or 100 µM 3iPKC by plot profiling. The results show that the ratio of inner to outer membrane signal intensity is significantly increased in single (1.25 ± 0.02 vs. 1.12 ± 0.01; *p* = 9.4×10-^9^; n = 26 and 23) and triple (1.32 ± 0.03 vs. 1.16 ± 0.01; *p* = 3.5×10?^7^; n = 20 and 24) inhibitor-treated groups compared with the controls (Figs. 4B–D and S4F). The outer membrane signal intensity of single (158.1 ± 3.6 vs. 160.2 ± 2.2; *p* = 0.63; n = 26 and 23) and triple (154.8 ± 3.6 vs. 160.3 ± 4.2; *p* = 0.34; n = 20 and 24) inhibitor-treated groups did not differ from that of the controls (Figs. 4E and S4G).

**Fig. 4.**
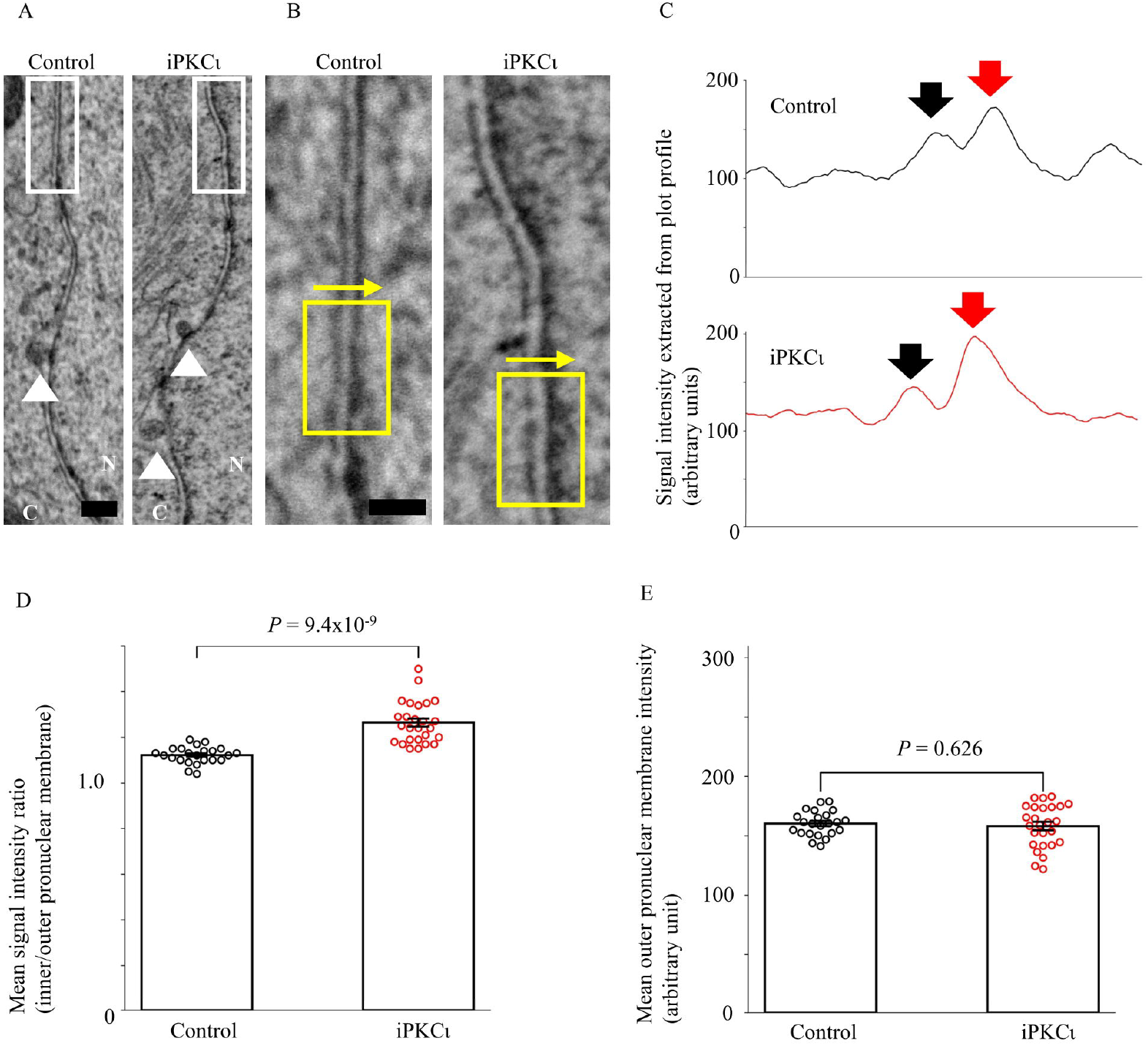
Effect of PKC inhibition on pronuclear membrane electron density. (A) Electron microscopy images (30,000×) of treated (iPKCι) or untreated mouse zygote pronuclear membranes. Treatment lasted 6 h, starting 3 h after IVF initiation. C: cytoplasm, N: pronucleus. White arrowheads indicate perinuclear vesicles/protrusions. (B) Enlarged views of white squares in A. Yellow squares mark regions used for plot profiling along yellow arrows. (C) Plot profiles of the signal intensity from B for both groups. Outer and inner nuclear membrane peaks are marked by black and red arrows, respectively. (D) Average ratio of inner–outer nuclear membrane peak intensities from plot profiles in C. Between 3 and 48 regions of the nuclear envelope lipid bilayer were analyzed per zygote. Ratios were averaged and compared between groups. (E) Average outer nuclear membrane peak intensity for both groups. Statistical analysis was carried out using two-tailed Student’s t-tests (*p* < 0.05: significantly different). IVF: *in vitro* fertilization, iPKCι: PKC inhibitor 1.

### 3.5. Perinuclear vesicles in 1-cell embryos produced by the calcium-independent method

To investigate the role of oocyte activation-associated cytoplasmic calcium transients in the formation of perinuclear vesicles in mouse 1-cell embryos, metaphase II (MII) oocytes were activated using TPEN that induces oocyte activation in a calcium-independent manner [22,33]. The resulting embryos were analyzed by electron microscopy. Nine h after the initiation of the TPEN treatment (TPEN 9 h), perinuclear vesicles were observed in all examined 1-cell embryos (n = 10, Figs. 2D, H, and L).

## 4. Discussion

Pharmacological inhibition observed in this study suggests that specific PKC isoforms are essential for mouse zygotes to reach the 2-cell stage. At 33 µM, iPKCι completely blocked this transition (0%), whereas Go 6983 had no effect (100%). At 10 µM, iPKCι moderately reduced blastocyst formation (61.8%), while Go 6983 fully inhibited it (0%; Fig. 1). PKCδ and PKCζ, targets of Go 6983, have been linked to blastocoel formation during blastocyst development [20], partially supporting these results. These findings suggest stage-specific effects of each PKC inhibitor during early development. Among PKC isoforms, PKCι is essential for embryogenesis in the knockout model [34], although its role in preimplantation stages remains unclear. The inhibitors used here lack high isoform specificity, which may help account for functional redundancy among PKCs [35]. Further studies are needed to pinpoint the isoforms critical for 2-cell development. Notably, mouse embryos can develop without fertilization-associated calcium transients [22], implicating calcium-independent PKC isoforms or alternative mechanisms. Based on the combination with prior evidence, that is, iPKCι as an optimized PKCι inhibitor [23] and PKCι’s predominant expression in mouse oocytes [11], PKCι emerges as a prime candidate for further investigation.

Electron microscopy showed that PKC inhibitor-treated mouse zygotes had (1) fewer perinuclear vesicles per unit pronuclear membrane circumference, (2) a reduced total vesicle area per unit length, and (3) increased electron density along the inner pronuclear membrane. These findings suggest that PKC signaling promotes perinuclear vesicle formation. The enhanced electron density may reflect an altered distribution of structural components or cargo accumulation normally cleared via perinuclear vesicles [2]. As the molecular basis of eeNEB in fertilized eggs remains poorly understood, further studies are required to elucidate its mechanisms and potential role in developmental progression.

Consistent with a previous report [7], perinuclear vesicles were observed in zygotes but not in GV oocytes or late 2-cell embryos. Vesicles emerged 3 h after IVF initiation, suggesting early onset of eeNEB post-fertilization. Similar perinuclear vesicles were also identified in TPEN-activated 1-cell embryos, indicating that their formation can occur independently of calcium signaling linked to oocyte activation. The correlations between NEB and eeNEB as well as between NEB-like processes across cell types remain unclear [8]. However, PKC inhibition reduces perinuclear vesicles, supporting the idea that eeNEB may involve PKC-dependent mechanisms, as proposed for NEB [2].

## 5. CRediT authorship contribution statement

Toru Suzuki: Conceptualization, Investigation, Formal analysis, Methodology, Resources, Funding acquisition, Project administration, Supervision, Visualization, Writing – original draft, Writing – review and editing. Yuriko Sakamaki: Investigation, Methodology, Resources, Writing – original draft.

## 8. Acknowledgements

We gratefully acknowledge the Stem Cell Laboratory at the Institute of Science Tokyo for their assistance with cell imaging.

## Glossary

BSA: Bovine serum albumin
COCs: Cumulus–oocyte complexes
DAG: Diacylglycerol
DMSO: Dimethyl sulfoxide
eeNEB: Early embryonic nuclear envelope budding
ELVAs: Endolysosomal vesicular assemblies
GV: Germinal vesicle
hCG: Human chorionic gonadotropin
HTF: Human tubal fluid
IBMX: 1-Methyl-3-Isobutylxanthine
IC_50_: Half maximal inhibitory concentration
IP_3_: Inositol 1,4,5-trisphosphate
IVF: *In vitro* fertilization
Ki: Inhibition constant
KSOM: Potassium simplex optimization medium
NEB: Nuclear envelope budding
NPC: Nuclear pore complex
PB: Phosphate buffer
PIP_2_: Phosphatidylinositol 4,5-bisphosphate
PKC: Protein kinase C
PLC: hospholipase C
TPEN: N,N,N′,N′-Tetrakis(2-pyridylmethyl)ethylenediamine
ZnSO_4_: Zinc sulfate

