## Supplemental Figure 1-6 for "Protein kinase C inhibitor suppresses 2-cell stage development and perinuclear vesicle formation in mouse zygotes"

**Supplemental Figure captions**

**Fig. S1.**

Bright-field images of embryos treated with individual PKC inhibitors**.** (A–B) Mouse zygotes generated via *in vitro* fertilization (IVF) were cultured for 1 (A) or 5 (B) days in the presence of the indicated PKC inhibitor (iPKCι, Go 6983, or sotrastaurin) at concentrations ranging from 100 to 0 μM (control). PKC: protein kinase C, iPKCι: PKCι inhibitor 1.

**Fig. S2.**

Bright-field images of embryos treated with multiple PKC inhibitors. (A–B) Mouse zygotes generated via *in vitro* fertilization (IVF) were cultured *in vitro* for 1 (A) or 5(B) days with either dual (Go 6983 + sotrastaurin; 2iPKC) or triple (iPKCι + Go 6983 + sotrastaurin; 3iPKC) PKC inhibitor combinations at concentrations ranging from 100 to 0 μM (control). PKC: protein kinase C, iPKCι: PKCι inhibitor 1.

**Fig. S3.**

Electron microscope images of mouse oocytes and embryos. Mouse germinal vesicle stage oocytes (GV oocyte) and 1- (IVF 3h) to 2-cell (IVF 30h) stage embryos produced by IVF were photographed using an electron microscope. (A–C) Cell nuclei at each stage. (D–K) Images of GV oocytes (D, E), IVF 3h (F–J), and IVF 30h (K). Black arrows, white arrows, white arrowheads, black arrowheads, and yellow arrowheads show endolysosomal vesicular assemblies (ELVAs), intranuclear annulate lamellae, cytoplasmic lattices, and electro-dance perinuclear microprotrusions, and a section of sperm tail detected in the same zygote in Fig. 2J, respectively. Black lines represent 5 (A, C), 2 (B), 1 (D, E, G), 0.5 (F, J, K) and 0.2 (H, I) μm. IVF: *in vitro* fertilization; ELVAs: endolysosomal vesicular assemblies.

**Fig. S4.**

Effects of PKC inhibitor mixture on perinuclear vesicle formation in mouse zygotes. (A) Representative brightfield images (200×) of embryos treated with 100 μM triple (iPKCι + Go 6983 + sotrastaurin; 3iPKC) PKC inhibitor combinations or untreated controls at 9 and 24 h after IVF initiation. Zygotes treated with 3iPKC arrest in the 1-cell stage 24 h after IVF initiation, while controls develop to the 2-cell stage. Black arrowheads mark second polar bodies. All images share the same scale; scale bar = 50 µm. (B) Electron microscopy image (1000×) of a pronucleus from a 3iPKC-treated zygote 9 h after IVF initiation, with an approximate formula for pronuclear perimeter calculation. The inset shows full zygote views. (C) Average number of perinuclear vesicles per unit pronuclear perimeter length. Values were averaged per zygote and then compared between treated (n = 24) and control (n = 22) groups. (D) Comparison of the average total perinuclear vesicle area per unit pronuclear perimeter length of 3iPKC-treated (n = 24) and control (n = 22) groups. (E) Comparison of the size of individual perinuclear vesicles of 3iPKC-treated (n = 140) and control (n = 41) groups. (F) Average ratio of inner–outer nuclear membrane peak intensities from plot profiles. Between 2 and 36 regions of the nuclear envelope lipid bilayer were analyzed per zygote. Ratios were averaged per zygote and compared between 3iPKC-treated (n = 24) and control (n = 20) groups. (G) Average outer nuclear membrane peak intensity from plot profiles for 3iPKC-treated (n = 24) and control (n = 20) groups. Statistical analysis was carried out via two-tailed Student's t-tests ($p < 0.05$: significantly different). IVF: *in vitro* fertilization.

**Fig. S5.**

Additional electron microscopy images focusing on pronuclear membranes of zygotes treated with or without iPKCι. Supplementary images corresponding to Fig. 4A are presented, showing zygotes cultured with (lower panels) or without (upper panels) 33 µM iPKCι. White arrowheads indicate the pronuclear membrane. In all panels, the pronucleus and cytoplasm are located on the right and left side, respectively, across the pronuclear membrane. The white scale bar in the bottom-right panel represents 0.2 µm and applies to all panels.

**Fig. S6.**

Additional electron microscopy images focusing on pronuclear membranes of zygotes treated with or without triple PKC inhibitors. Supplementary examples corresponding to Fig. S4F are shown, displaying zygotes cultured with (lower panels, 3iPKC) or without (upper panels, control) a 100 µM 3 PKC inhibitor cocktail (iPKCι + Go 6983 + sotrastaurin). White arrowheads indicate the pronuclear membrane. In all panels, the pronucleus and cytoplasm are located on the right and left side, respectively, across the pronuclear membrane. The white scale bar in the bottom-right panel represents 0.2 µm and applies to all panels.

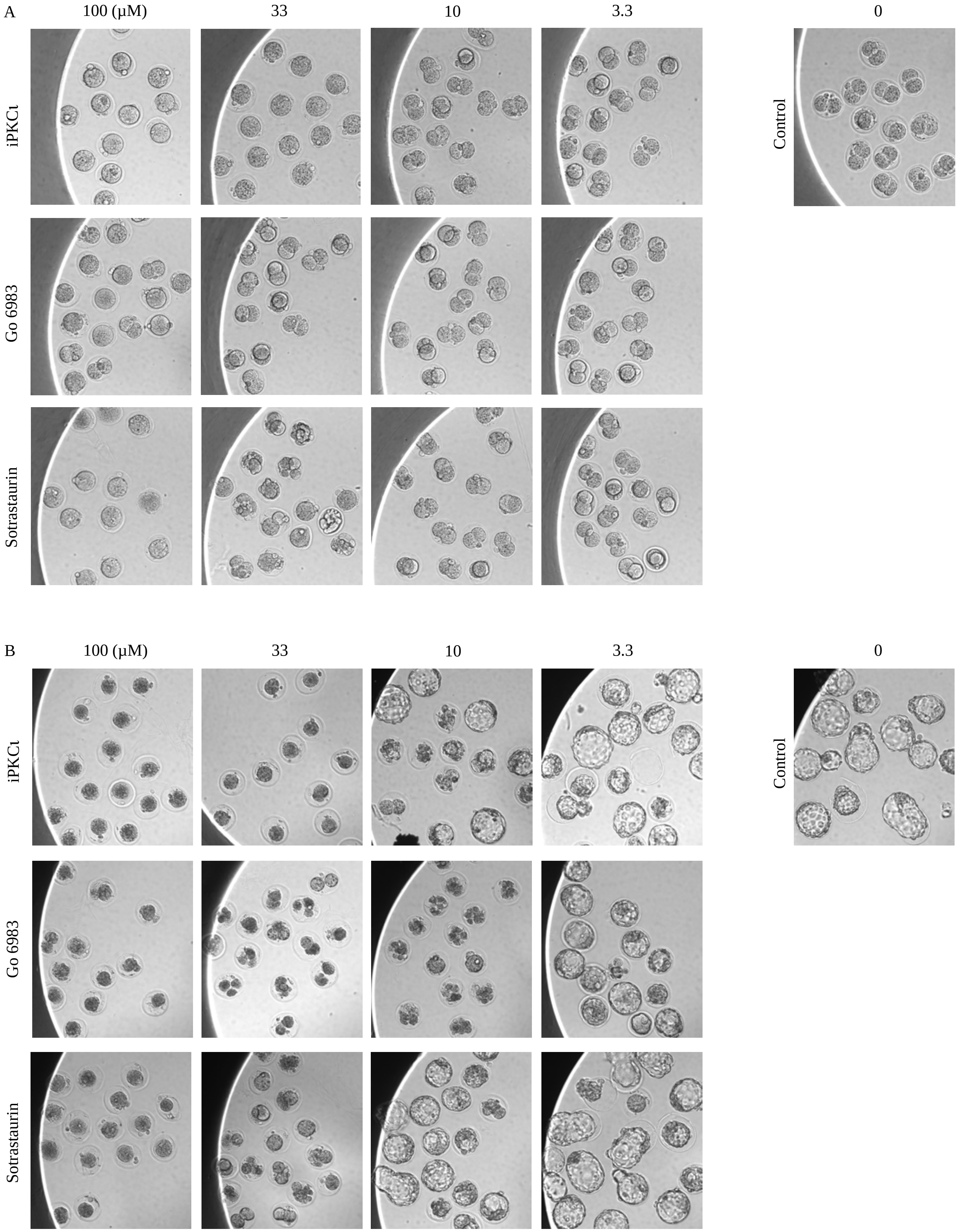
A
100 (μM)
33
10
3.3
0
iPKCι
Go 6983
Sotrastaurin
Control
B
100 (μM)
33
10
3.3
0
iPKCι
Go 6983
Sotrastaurin
Control

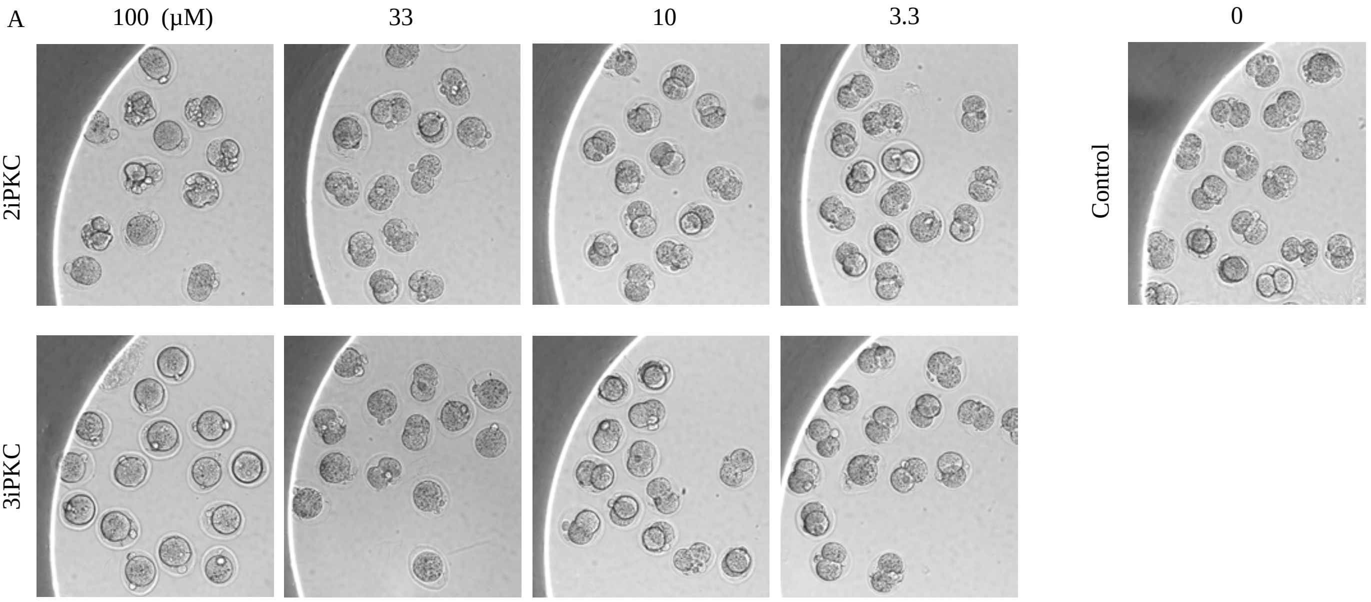
A
100 (μM)
33
10
3.3
0
2iPKC
3iPKC
Control

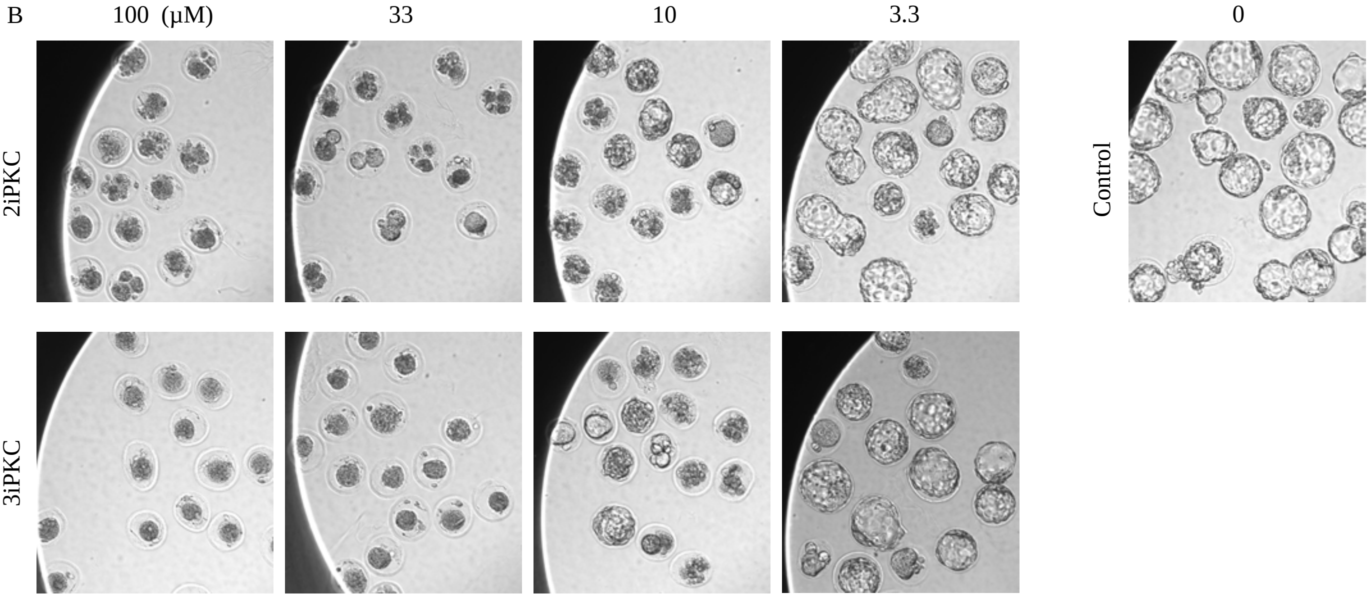
B
100 (μM)
33
10
3.3
0
2iPKC
3iPKC
Control

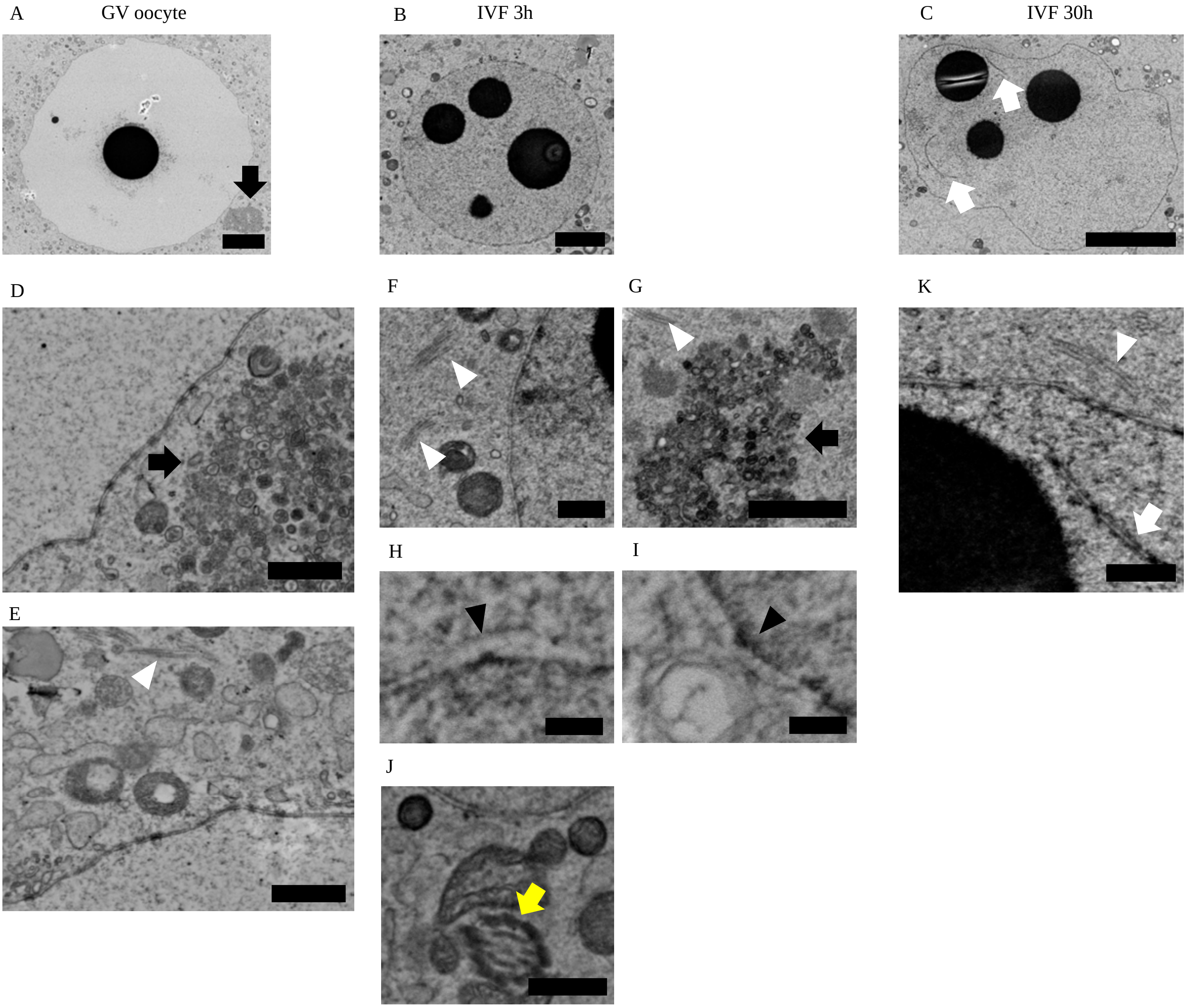
A
GV oocyte
B
IVF 3h
C
IVF 30h
D
E
F
G
H
I
J
K

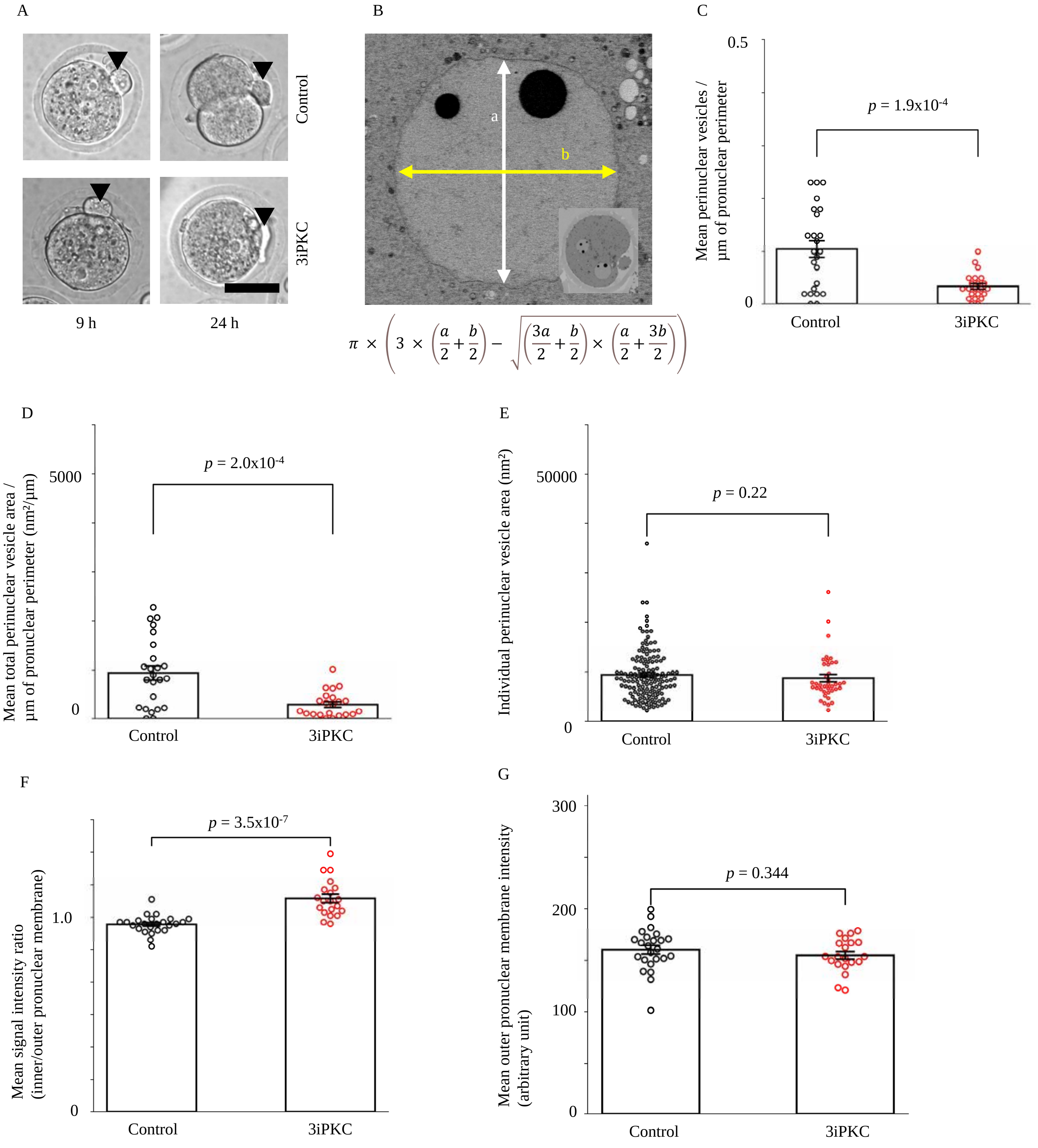
A
Control
3iPKC
9 h
24 h
B
a
b
π × (3 × (a/2 + b/2) − √((3a/2 + b/2) × (a/2 + 3b/2)))
C
Mean perinuclear vesicles /
μm of pronuclear perimeter
0.5
0
p = 1.9x10-4
Control
3iPKC
D
Mean total perinuclear vesicle area /
μm of pronuclear perimeter (nm²/μm)
5000
0
p = 2.0x10-4
Control
3iPKC
E
Individual perinuclear vesicle area (nm²)
50000
0
p = 0.22
Control
3iPKC
F
Mean signal intensity ratio
(inner/outer pronuclear membrane)
1.0
0
p = 3.5x10-7
Control
3iPKC
G
Mean outer pronuclear membrane intensity
(arbitrary unit)
300
200
100
0
p = 0.344
Control
3iPKC

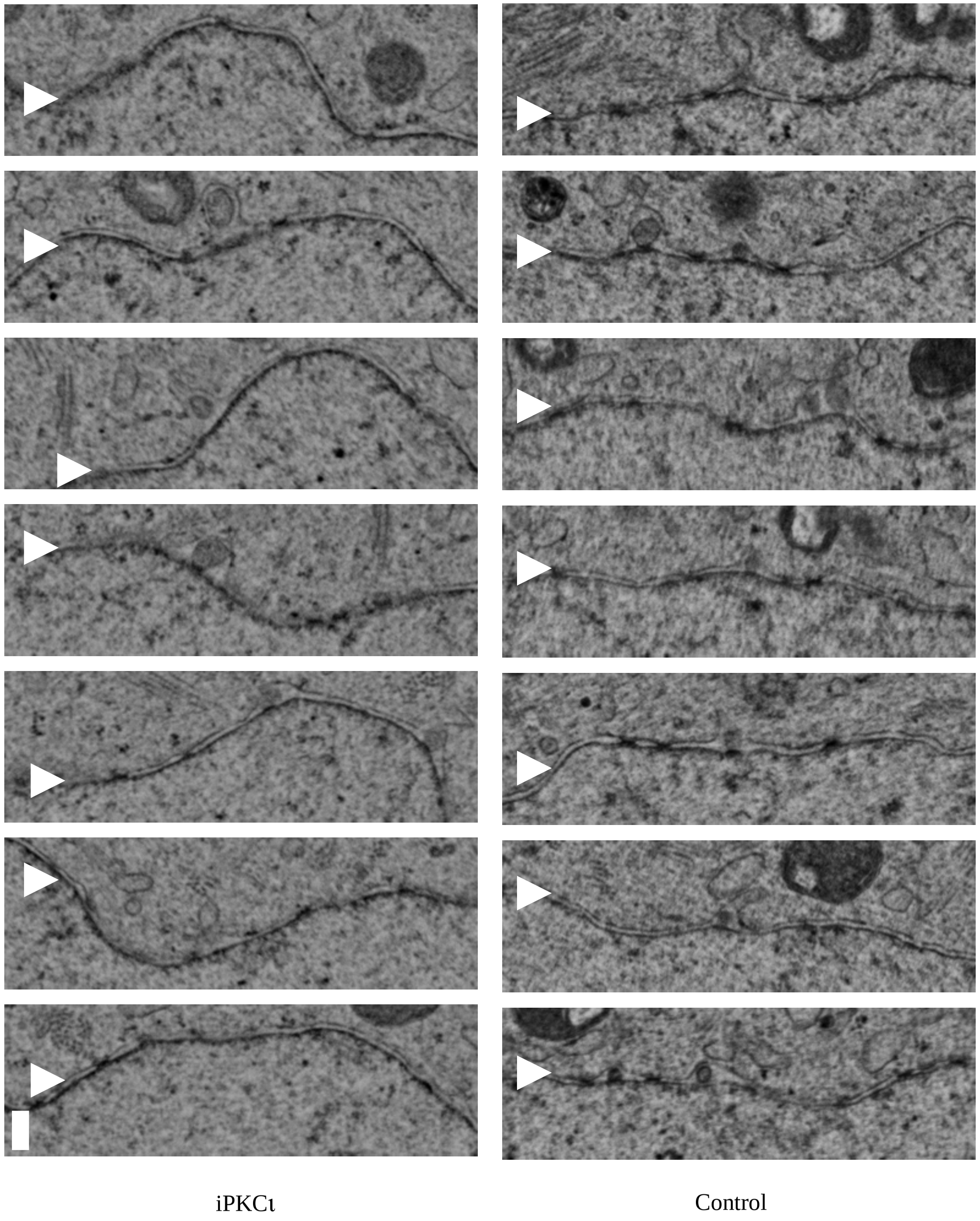
iPKCι
Control

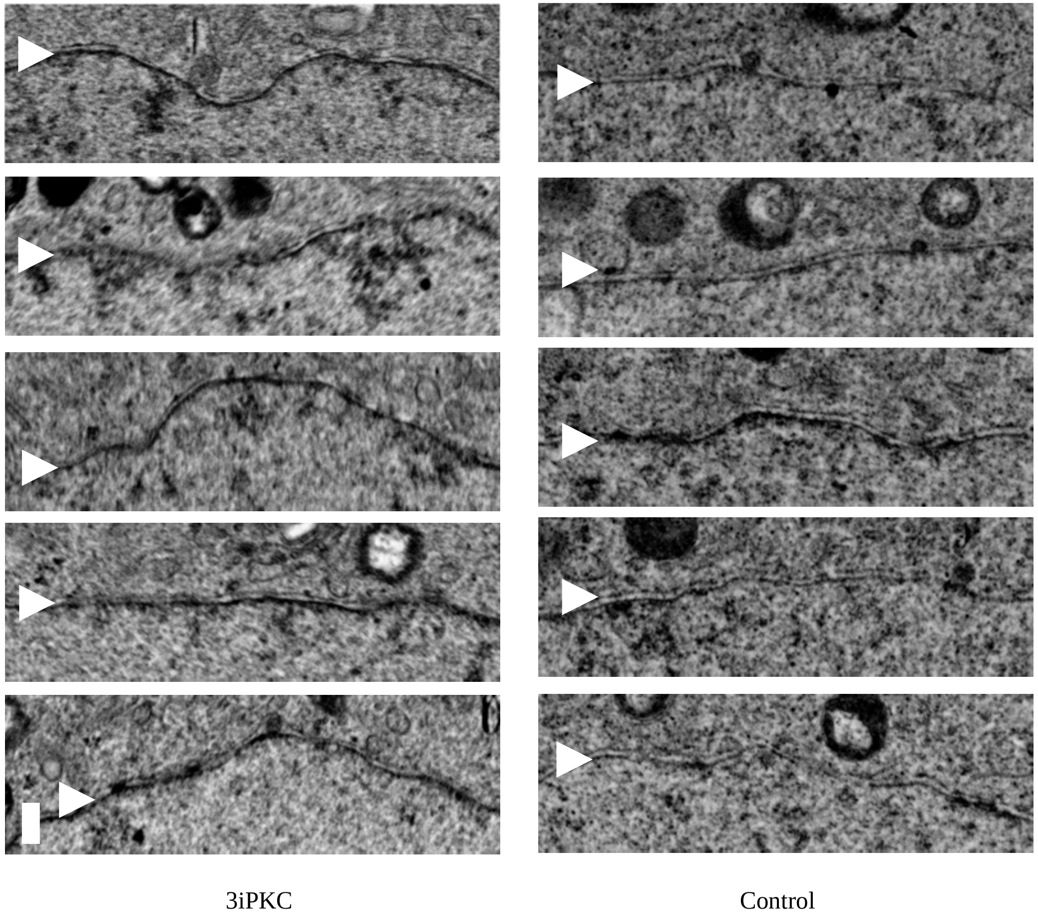
3iPKC
Control
