## Supplemental Table 1-4 for "Protein kinase C inhibitor suppresses 2-cell stage development and perinuclear vesicle formation in mouse zygotes"

Supplemental Table 1. Preimplantation development of mouse embryos cultured with or without single PKC inhibitor.

| μM | 100 | | | 33 | | | 10 | | | 3.3 | | | 0 |
|---|---|---|---|---|---|---|---|---|---|---|---|---|---|
| **Chemical** | **iPKCι** | **Go 6983** | **sotrastaurin** | **iPKCι** | **Go 6983** | **sotrastaurin** | **iPKCι** | **Go 6983** | **sotrastaurin** | **iPKCι** | **Go 6983** | **sotrastaurin** | **control** |
| zygote (n) | 34 | 34 | 34 | 34 | 34 | 34 | 34 | 34 | 34 | 34 | 31 | 34 | 32 |
| 2-cell (n) | 0 | 7 | 0 | 0 | 34 | 14 | 32 | 34 | 32 | 31 | 29 | 32 | 31 |
| (% ± SD)*[1] | (0.0 ± 0.0) | (20.6 ± 8.8) | (0.0 ± 0.0) | (0.0 ± 0.0) | (100 ± 0.0) | (41.2 ± 5.9) | (94.1 ± 5.9) | (100 ± 0.0) | (94.1 ± 0.0) | (91.2 ± 2.9) | (93.5 ± 0.6) | (94.1 ± 5.9) | (93.9 ± 0.2) |
| 4-cell (n) | 0 | 0 | 0 | 0 | 3 | 0 | 27 | 32 | 32 | 31 | 28 | 30 | 30 |
| (% ± SD)*[1] | (0.0 ± 0.0) | (0.0 ± 0.0) | (0.0 ± 0.0) | (0.0 ± 0.0) | (8.8 ± 8.8) | (0.0 ± 0.0) | (79.4 ± 8.8) | (94.1 ± 0.0) | (94.1 ± 0.0) | (91.2 ± 2.9) | (90.5 ± 2.3) | (88.2 ± 0.0) | (90.8 ± 3.3) |
| morula (n) | 0 | 0 | 0 | 0 | 0 | 0 | 27 | 15 | 32 | 31 | 28 | 30 | 29 |
| (% ± SD)*[1] | (0.0 ± 0.0) | (0.0 ± 0.0) | (0.0 ± 0.0) | (0.0 ± 0.0) | (0.0 ± 0.0) | (0.0 ± 0.0) | (79.4 ± 8.8) | (44.1 ± 8.8) | (94.1 ± 0.0) | (91.2 ± 2.9) | (90.5 ± 2.3) | (88.2 ± 0.0) | (87.9 ± 0.4) |
| blastocyst (n) | 0 | 0 | 0 | 0 | 0 | 0 | 21 | 0 | 31 | 28 | 25 | 30 | 28 |
| (% ± SD)*[1] | (0.0 ± 0.0) | (0.0 ± 0.0) | (0.0 ± 0.0) | (0.0 ± 0.0) | (0.0 ± 0.0) | (0.0 ± 0.0) | (61.8 ± 2.9) | (0.0 ± 0.0) | (91.2 ± 2.9) | (82.4 ± 11.8) | (81.1 ± 4.6) | (88.2 ± 0.0) | (84.7 ± 3.5) |

The number and percentage (mean ± standard deviation) of mouse embryos that reached each specified developmental stage following culture with or without a single PKC inhibitor at the indicated concentrations are shown.
Data was collected from 2 replecates. n: numbers.
*[1]% (± SD) of total number of zygotes.
PKC: protein kinase C, iPKCι: PKCι inhibitor 1.

Supplemental Table 2. *p* -values for all of comparison groups in preimplantation development of mouse embryos cultured with or without single PKC inhibitor at the indicated concentration.

| | 100μM | | | 33μM | | | 10μM | | | 3.3μM | | |
|---|---|---|---|---|---|---|---|---|---|---|---|---|
| 2-cell | **Comparison group** | **Raw p-value** | **Holm-adjusted p-value** | **Comparison group** | **Raw p-value** | **Holm-adjusted p-value** | **Comparison group** | **Raw p-value** | **Holm-adjusted p-value** | **Comparison group** | **Raw p-value** | **Holm-adjusted p-value** |
| | iPKCι vs control | 4.99E-18 | 2.99E-17 | iPKCι vs Go 6983 | 7.03E-20 | 4.22E-19 | Go 6983 vs control | 0.4848 | 1 | Go 6983 vs control | 0.6128 | 1 |
| | Sotrastaurin vs control | 4.99E-18 | 2.99E-17 | iPKCι vs control | 4.99E-18 | 2.50E-17 | iPKCι vs Go 6983 | 0.4925 | 1 | iPKCι vs control | 0.6139 | 1 |
| | Go 6983 vs control | 5.28E-11 | 2.11E-10 | Go 6983 vs Sotrastaurin | 3.39E-08 | 1.36E-07 | Go 6983 vs Sotrastaurin | 0.4925 | 1 | iPKCι vs Go 6983 | 1 | 1 |
| | iPKCι vs Go 6983 | 1.11E-02 | 3.33E-02 | Sotrastaurin vs control | 5.98E-07 | 1.79E-06 | iPKCι vs Sotrastaurin | 1 | 1 | iPKCι vs Sotrastaurin | 1 | 1 |
| | Go 6983 vs Sotrastaurin | 1.11E-02 | 3.33E-02 | iPKCι vs Sotrastaurin | 2.26E-05 | 4.52E-05 | iPKCι vs control | 1 | 1 | Go 6983 vs Sotrastaurin | 1 | 1 |
| | iPKCι vs Sotrastaurin | 1 | 1 | Go 6983 vs control | 0.485 | 0.485 | Sotrastaurin vs control | 1 | 1 | Sotrastaurin vs control | 1 | 1 |

| | 100μM | | | 33μM | | | 10μM | | | 3.3μM | | |
|---|---|---|---|---|---|---|---|---|---|---|---|---|
| 4-cell | **Comparison group** | **Raw p-value** | **Holm-adjusted p-value** | **Comparison group** | **Raw p-value** | **Holm-adjusted p-value** | **Comparison group** | **Raw p-value** | **Holm-adjusted p-value** | **Comparison group** | **Raw p-value** | **Holm-adjusted p-value** |
| | iPKCι vs control | 8.99E-17 | 5.395E-16 | iPKCι vs control | 8.991E-17 | 5.395E-16 | iPKCι vs Go 6983 | 0.1497 | 0.898 | Go 6983 vs control | 0.6719 | 1 |
| | Go 6983 vs control | 8.991E-17 | 5.395E-16 | Sotrastaurin vs control | 8.991E-17 | 5.395E-16 | iPKCι vs Sotrastaurin | 0.1497 | 0.898 | Sotrastaurin vs control | 0.6733 | 1 |
| | Sotrastaurin vs control | 8.991E-17 | 5.395E-16 | Go 6983 vs control | 8.272E-13 | 3.309E-12 | iPKCι vs control | 0.1506 | 0.898 | iPKCι vs Go 6983 | 1 | 1 |
| | iPKCι vs Go 6983 | 1 | 1 | iPKCι vs Go 6983 | 0.2388 | 0.7164 | Go 6983 vs Sotrastaurin | 1 | 1 | iPKCι vs Sotrastaurin | 1 | 1 |
| | iPKCι vs Sotrastaurin | 1 | 1 | Go 6983 vs Sotrastaurin | 0.2388 | 0.7164 | Go 6983 vs control | 1 | 1 | iPKCι vs control | 1 | 1 |
| | Go 6983 vs Sotrastaurin | 1 | 1 | iPKCι vs Sotrastaurin | 1 | 1 | Sotrastaurin vs control | 1 | 1 | Go 6983 vs Sotrastaurin | 1 | 1 |

| | 100μM | | | 33μM | | | 10μM | | | 3.3μM | | |
|---|---|---|---|---|---|---|---|---|---|---|---|---|
| morula | **Comparison group** | **Raw p-value** | **Holm-adjusted p-value** | **Comparison group** | **Raw p-value** | **Holm-adjusted p-value** | **Comparison group** | **Raw p-value** | **Holm-adjusted p-value** | **Comparison group** | **Raw p-value** | **Holm-adjusted p-value** |
| | iPKCι vs control | 1.109E-15 | 6.653E-15 | iPKCι vs control | 1.109E-15 | 6.653E-15 | Go 6983 vs Sotrastaurin | 0.00001161 | 0.00006965 | iPKCι vs Go 6983 | 1 | 1 |
| | Go 6983 vs control | 1.109E-15 | 6.653E-15 | Go 6983 vs control | 1.109E-15 | 6.653E-15 | Go 6983 vs control | 0.00006661 | 0.0003331 | iPKCι vs Sotrastaurin | 1 | 1 |
| | Sotrastaurin vs control | 1.109E-15 | 6.653E-15 | Sotrastaurin vs control | 1.109E-15 | 6.653E-15 | iPKCι vs Go 6983 | 0.005548 | 0.02219 | iPKCι vs control | 1 | 1 |
| | iPKCι vs Go 6983 | 1 | 1 | iPKCι vs Go 6983 | 1 | 1 | iPKCι vs Sotrastaurin | 0.1497 | 0.449 | Go 6983 vs Sotrastaurin | 1 | 1 |
| | iPKCι vs Sotrastaurin | 1 | 1 | iPKCι vs Sotrastaurin | 1 | 1 | iPKCι vs control | 0.306 | 0.612 | Go 6983 vs control | 1 | 1 |
| | Go 6983 vs Sotrastaurin | 1 | 1 | Go 6983 vs Sotrastaurin | 1 | 1 | Sotrastaurin vs control | 0.6679 | 0.6679 | Sotrastaurin vs control | 1 | 1 |

| | 100μM | | | 33μM | | | 10μM | | | 3.3μM | | |
|---|---|---|---|---|---|---|---|---|---|---|---|---|
| blastocyst | **Comparison group** | **Raw p-value** | **Holm-adjusted p-value** | **Comparison group** | **Raw p-value** | **Holm-adjusted p-value** | **Comparison group** | **Raw p-value** | **Holm-adjusted p-value** | **Comparison group** | **Raw p-value** | **Holm-adjusted p-value** |
| | iPKCι vs control | 1.053E-14 | 6.321E-14 | iPKCι vs control | 1.053E-14 | 6.321E-14 | Go 6983 vs Sotrastaurin | 5.462E-16 | 3.277E-15 | Go 6983 vs Sotrastaurin | 0.4995 | 1 |
| | Go 6983 vs control | 1.053E-14 | 6.321E-14 | Go 6983 vs control | 1.053E-14 | 6.321E-14 | Go 6983 vs control | 1.053E-14 | 5.267E-14 | Go 6983 vs control | 0.5092 | 1 |
| | Sotrastaurin vs control | 1.053E-14 | 6.321E-14 | Sotrastaurin vs control | 1.053E-14 | 6.321E-14 | iPKCι vs Go 6983 | 9.888E-09 | 3.955E-08 | iPKCι vs Sotrastaurin | 0.7337 | 1 |
| | iPKCι vs Go 6983 | 1 | 1 | iPKCι vs Go 6983 | 1 | 1 | iPKCι vs Sotrastaurin | 0.008709 | 0.02613 | iPKCι vs control | 0.7344 | 1 |
| | iPKCι vs Sotrastaurin | 1 | 1 | iPKCι vs Sotrastaurin | 1 | 1 | iPKCι vs control | 0.02406 | 0.04812 | iPKCι vs Go 6983 | 1 | 1 |
| | Go 6983 vs Sotrastaurin | 1 | 1 | Go 6983 vs Sotrastaurin | 1 | 1 | Sotrastaurin vs control | 0.7046 | 0.7046 | Sotrastaurin vs control | 1 | 1 |

The number of embryos that developed to each specified stage (Supplemental Table 1) was analyzed using Fisher's exact test for all pairwise comparisons, with p-values adjusted using the Holm method.
PKC: protein kinase C, iPKCι: PKCι inhibitor 1.

Supplemental Table 3. Preimplantation development of mouse embryos cultured with or without double or triple PKC inhibitors.

| μM (total) | 100 | | 33 | | 10 | | 3.3 | | 0 |
|---|---|---|---|---|---|---|---|---|---|
| **Chemical** | **3iPKC** | **2iPKC** | **3iPKC** | **2iPKC** | **3iPKC** | **2iPKC** | **3iPKC** | **2iPKC** | **control** |
| zygote (n) | 32 | 32 | 32 | 32 | 32 | 32 | 32 | 32 | 32 |
| 2-cell (n) | 0 | 0 | 3 | 26 | 30 | 30 | 28 | 30 | 30 |
| (% ± SD)*[1] | (0.0 ± 0.0) | (0.0 ± 0.0) | (9.4± 3.1) | (81.3 ± 0.0) | (93.8 ± 6.3) | (93.8 ± 0.0) | (87.5 ± 6.3) | (93.8 ± 0.0) | (93.8 ± 6.3) |
| 4-cell (n) | 0 | 0 | 0 | 2 | 23 | 27 | 23 | 29 | 27 |
| (% ± SD)*[1] | (0.0 ± 0.0) | (0.0 ± 0.0) | (0.0 ± 0.0) | (6.3 ± 0.0) | (71.9 ± 3.1) | (84.4 ± 9.4) | (71.9 ± 9.4) | (90.6 ± 3.1) | (84.4 ± 3.1) |
| morula (n) | 0 | 0 | 0 | 0 | 21 | 26 | 23 | 29 | 27 |
| (% ± SD)*[1] | (0.0 ± 0.0) | (0.0 ± 0.0) | (0.0 ± 0.0) | (0.0 ± 0.0) | (65.6 ± 9.4) | (81.3 ± 6.3) | (71.9 ± 9.4) | (90.6 ± 3.1) | (84.4 ± 3.1) |
| blastocyst (n) | 0 | 0 | 0 | 0 | 10 | 11 | 21 | 27 | 26 |
| (% ± SD)*[1] | (0.0 ± 0.0) | (0.0 ± 0.0) | (0.0 ± 0.0) | (0.0 ± 0.0) | (31.3 ± 25.0) | (34.4 ± 3.1) | (65.6 ± 9.4) | (84.4 ± 3.1) | (81.3 ± 0.0) |

The number and percentage (mean ± standard deviation) of mouse embryos that reached each specified developmental stage after being cultured with or without double (2iPKC: Go 6983 + sotrastaurin) or triple (3iPKC: iPKCι + Go 6983 + sotrastaurin) PKC inhibitors at the indicated concentrations are shown.
Data was collected from 2 replecates. n: numbers.
*[1]% (± SD) of total number of zygotes.
SD: standard diviation.

Supplemental table 4. *p* -values for all of comparison groups in preimplantation development of mouse embryos cultured with or without double pr triple PKC inhibitors at the indicated total concentration.

| | 100μM | | | 33μM | | | 10μM | | | 3.3μM | | |
|---|---|---|---|---|---|---|---|---|---|---|---|---|
| 2-cell | **Comparison group** | **Raw p-value** | **Holm-adjusted p-value** | **Comparison group** | **Raw p-value** | **Holm-adjusted p-value** | **Comparison group** | **Raw p-value** | **Holm-adjusted p-value** | **Comparison group** | **Raw p-value** | **Holm-adjusted p-value** |
| | 3iPKC vs control | 6.12E-16 | 1.84E-15 | 3iPKC vs control | 2.79E-12 | 8.36E-12 | 3iPKC vs 2iPKC | 1 | 1 | 3iPKC vs 2iPKC | 0.671867 | 1 |
| | 2iPKC vs control | 6.12E-16 | 1.84E-15 | 3iPKC vs 2iPKC | 6.62E-09 | 1.32E-08 | 3iPKC vs control | 1 | 1 | 3iPKC vs control | 0.671867 | 1 |
| | 3iPKC vs 2iPKC | 1 | 1 | 2iPKC vs control | 0.256519 | 0.256519 | 2iPKC vs control | 1 | 1 | 2iPKC vs control | 1 | 1 |

| | 100μM | | | 33μM | | | 10μM | | | 3.3μM | | |
|---|---|---|---|---|---|---|---|---|---|---|---|---|
| 4-cell | **Comparison group** | **Raw p-value** | **Holm-adjusted p-value** | **Comparison group** | **Raw p-value** | **Holm-adjusted p-value** | **Comparison group** | **Raw p-value** | **Holm-adjusted p-value** | **Comparison group** | **Raw p-value** | **Holm-adjusted p-value** |
| | 3iPKC vs control | 4.76E-13 | 1.43E-12 | 3iPKC vs control | 4.76E-13 | 1.43E-12 | 3iPKC vs 2iPKC | 0.36492 | 1 | 3iPKC vs 2iPKC | 0.106859 | 0.320578 |
| | 2iPKC vs control | 4.76E-13 | 1.43E-12 | 2iPKC vs control | 1.46E-10 | 2.91E-10 | 3iPKC vs control | 0.36492 | 1 | 3iPKC vs control | 0.36492 | 0.729841 |
| | 3iPKC vs 2iPKC | 1 | 1 | 3iPKC vs 2iPKC | 0.492063 | 0.492063 | 2iPKC vs control | 1 | 1 | 2iPKC vs control | 0.707846 | 0.729841 |

| | 100μM | | | 33μM | | | 10μM | | | 3.3μM | | |
|---|---|---|---|---|---|---|---|---|---|---|---|---|
| morula | **Comparison group** | **Raw p-value** | **Holm-adjusted p-value** | **Comparison group** | **Raw p-value** | **Holm-adjusted p-value** | **Comparison group** | **Raw p-value** | **Holm-adjusted p-value** | **Comparison group** | **Raw p-value** | **Holm-adjusted p-value** |
| | 3iPKC vs control | 4.76E-13 | 1.43E-12 | 3iPKC vs control | 4.76E-13 | 1.43E-12 | 3iPKC vs control | 0.147699 | 0.443097 | 3iPKC vs 2iPKC | 0.106859 | 0.320578 |
| | 2iPKC vs control | 4.76E-13 | 1.43E-12 | 2iPKC vs control | 4.76E-13 | 1.43E-12 | 3iPKC vs 2iPKC | 0.257394 | 0.514787 | 3iPKC vs control | 0.36492 | 0.729841 |
| | 3iPKC vs 2iPKC | 1 | 1 | 3iPKC vs 2iPKC | 1 | 1 | 2iPKC vs control | 1 | 1 | 2iPKC vs control | 0.707846 | 0.729841 |

| | 100μM | | | 33μM | | | 10μM | | | 3.3μM | | |
|---|---|---|---|---|---|---|---|---|---|---|---|---|
| blastocyst | **Comparison group** | **Raw p-value** | **Holm-adjusted p-value** | **Comparison group** | **Raw p-value** | **Holm-adjusted p-value** | **Comparison group** | **Raw p-value** | **Holm-adjusted p-value** | **Comparison group** | **Raw p-value** | **Holm-adjusted p-value** |
| | 3iPKC vs control | 3.01E-12 | 9.04E-12 | 3iPKC vs control | 3.01E-12 | 9.04E-12 | 3iPKC vs control | 0.000115313 | 0.000345938 | 3iPKC vs 2iPKC | 0.147699 | 0.443097 |
| | 2iPKC vs control | 3.01E-12 | 9.04E-12 | 2iPKC vs control | 3.01E-12 | 9.04E-12 | 2iPKC vs control | 0.0003094 | 0.0006188 | 3iPKC vs control | 0.257394 | 0.514787 |
| | 3iPKC vs 2iPKC | 1 | 1 | 3iPKC vs 2iPKC | 1 | 1 | 3iPKC vs 2iPKC | 1 | 1 | 2iPKC vs control | 1 | 1 |

The number of embryos that developed to each specified stage (Supplemental Table 3) was analyzed using Fisher’s exact test for all pairwise comparisons, with p-values adjusted using the Holm method.
The indicated concentrations represent the total concentration of the inhibitor mixture, in which each individual inhibitor is present at an equal concentration.
2iPKC: Go 6983 + sotrastaurin, 3iPKC: iPKCι + Go 6983 + sotrastaurin, PKC: protein kinase C, iPKCι: PKCι inhibitor 1.
